# Functional specialization of the gibberellin receptor GIBBERELLIN-INSENSITIVE DWARF 1C in plant neighbour detection

**DOI:** 10.64898/2026.07.09.737222

**Authors:** Putri Prasetyaningrum, Vincent Hofheinz Crisostomo, Matthias Reimers, Sebastian Krueger, Andreas Hiltbrunner

## Abstract

Plants detect neighbours through a reduced red-to-far-red ratio (R:FR), triggering elongation growth that reduces crop yield. Although Gibberellin (GA) is required for the neighbour-proximity (NP) elongation response, bioactive GA levels do not increase sufficiently to account for elongation magnitude, suggesting GA sensitivity as an additional regulated variable. Here, we show that GID1C, one of three Arabidopsis GA receptors, is the primary GA receptor involved in NP-induced elongation. GID1C protein accumulates selectively in hypocotyls and root tips under low R:FR without an increase in bioactive GA. The *gid1c* mutant shows a reduced elongation response that exogenous GA treatment cannot rescue. Transcriptome profiling reveals that GID1C controls 86% of the NP-responsive transcriptome, including genes for cell growth, division, and transcriptional regulation. Hub analysis identifies ICE1 as a GID1C-repressed transcriptional brake. *ICE1* transcript is suppressed under low R:FR in a GID1C-dependent manner, and a phosphorylation-resistant *ICE1* allele blocks NP-induced elongation. Together, these findings establish GA perception as an additional regulatory layer in NP, with subfunctionalisation among GID1 paralogs shaping the response to neighbouring plants.

## Introduction

Plants in dense vegetation detect neighbours through a reduced red-to-far-red light ratio (R:FR). Chlorophyll absorbs red light while surrounding foliage reflects far-red light. This signal is perceived by PHYTOCHROME B (PHYB), a dimeric photoreceptor active in full sunlight that represses PHYTOCHROME-INTERACTING FACTORS (PIFs). Low R:FR inactivates PHYB, thereby relieving repression of PIFs and triggering elongation in shade-intolerant plants, such as *Arabidopsis thaliana*^1^. Low R:FR arises in two ecologically distinct contexts^2^. Canopy shade combines reduced light intensity, depleted blue light, and low R:FR beneath overhead vegetation. Neighbour proximity (NP), which is the focus of this study, lowers R:FR alone, as lateral foliage reflects far-red onto plants still receiving full sunlight. The phytohormone Gibberellin (GA) is a central regulator of shade-induced elongation^3^, acting through degradation of the DELLA growth-repressor proteins^4^. GA is required for elongation under low R:FR since blocking its biosynthesis suppresses the response. However, bioactive GA levels do not increase sufficiently to account for the elongation under WL+FR^5–7^, implicating GA sensitivity as an additional regulated variable.

Canonical GA signalling starts with binding of bioactive GA to its receptor GIBBERELLIN INSENSITIVE DWARF1 (GID1). GID1 then recruits the GRAS-family DELLA growth-repressor proteins and binds the SCF–SLY1 complex (Skp1, Cullin, and the F-box protein SLEEPY1/SLY1), leading to ubiquitination and degradation of DELLA^8,9^. GA therefore promotes growth by relieving DELLA repression. Arabidopsis possesses three GA receptors, GID1A, GID1B and GID1C. Loss of all three causes complete GA insensitivity^10^, despite elevated bioactive GA level^11^. In contrast, GA-biosynthesis mutants have reduced GA levels, but are rescued by exogenous GA^12,13^. Whereas the role of GA in NP elongation has been studied extensively, the contribution of individual GID1 paralogs remains largely unexplored. The three paralogs are functionally not equivalent. GID1B binds DELLA independently of GA and more strongly than GID1A and GID1C, and each contributes distinctly to germination, fertility, and environmental robustness^14,15^.

To directly test GID1 function, we mimicked neighbour proximity using white light (WL) supplemented with far-red light (WL+FR) and combined phenotypic and transcriptomic profiling. In *Arabidopsis thaliana*, WL+FR selectively increases GID1C protein abundance in hypocotyls and root tips without altering GA levels, and *gid1c* is specifically deficient in NP-induced hypocotyl elongation. Factorial RNAseq data analysis shows GID1C alone drives 86% of the NP-response transcriptome, controlling genes for cell growth, division, and meristem function. This also includes the transcriptional hub *INDUCER OF CBF EXPRESSION 1* (*ICE1/SCREAM/SCRM*). These results reveal that GA receptor identity and abundance actively shape how plants reprogram growth in response to neighbours, uncovering a hidden layer of specificity in hormone signalling.

## Result

### GID1C is required for the full response to neighbour proximity

As NP is perceived through PHYB, we first confirmed that our WL+FR treatment acts through this pathway (Figure S1A). Downstream of PHYB, the roles of the individual GID1 receptors in NP remain unclear. We therefore examined hypocotyl elongation in GID1 single (Figure 1A) and double mutants (Figure S1B) under WL and WL+FR, which lowers R:FR to mimic NP. Among the single mutants^10^, *gid1a* resembled the wild type, *gid1b* had shorter hypocotyls regardless of light, and *gid1c* showed reduced elongation relative to Col-0 specifically under WL+FR (Figure 1A), revealing condition-specific roles for individual GID1s. To test whether GID1C is required for the GA-dependent NP response, we treated Col-0 and *gid1c* seedlings under WL+FR with exogenous GA_4+7_ or the GA-biosynthesis inhibitor paclobutrazol (PAC). In Col-0, elongation was suppressed by PAC and enhanced by GA, whereas *gid1c* elongated minimally even with exogenous GA (Figure 1B), indicating that GID1C is essential for GA sensitivity during the NP response. Reintroducing GID1C (*pGID1C:GID1C-mCitrine#4*) restored the WL+FR response (Figure 1C), confirming the defect is due to loss of GID1C.

**Figure 1.**
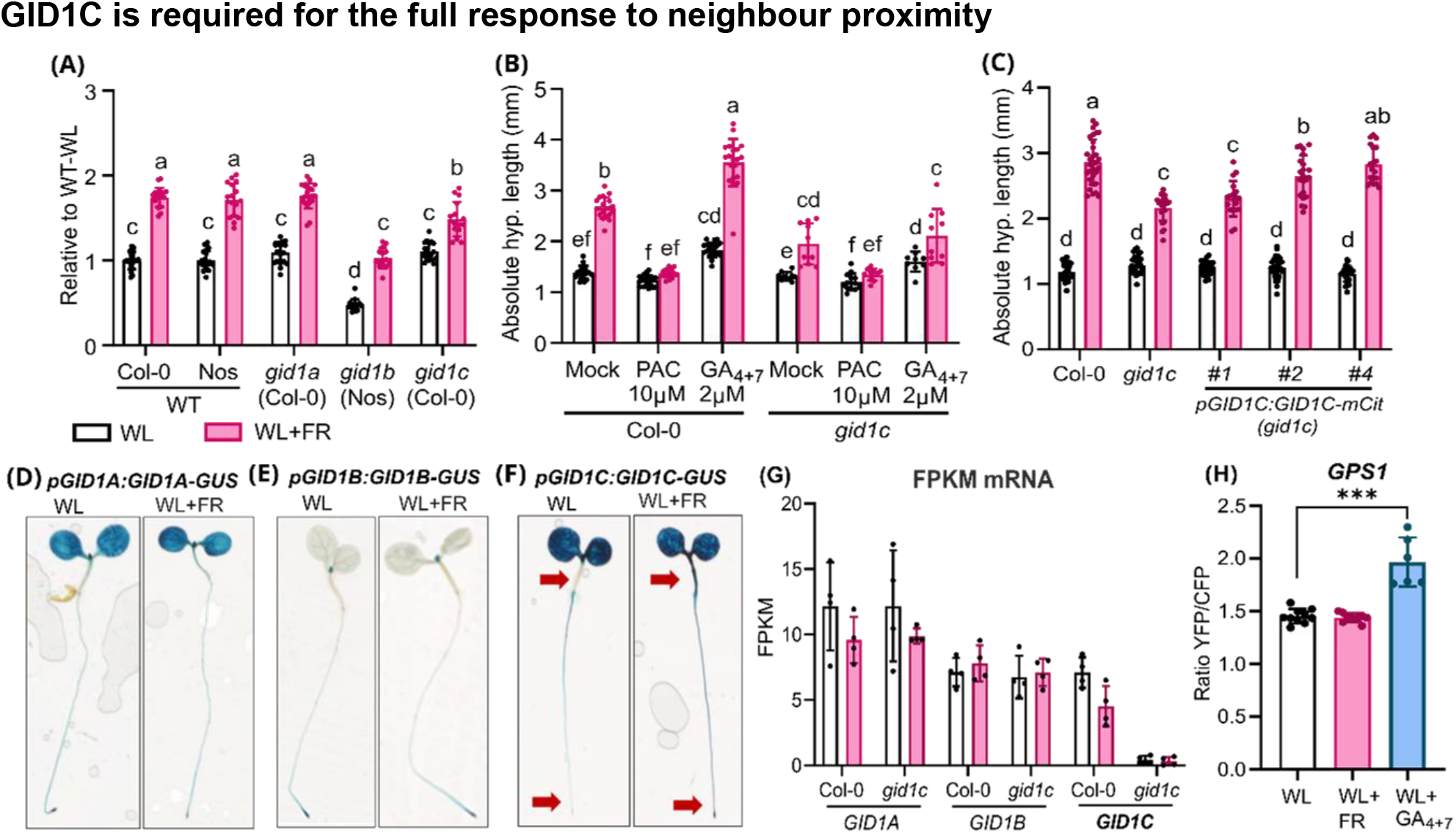
GID1C is required for hypocotyl elongation in shade avoidance. Hypocotyl length of **(A)** single mutants relative to their wildtype background indicated in brackets. **(B)** Absolute hypocotyl length of *gid1c* and **(C)** *pGID1C:GID1C-mCitrine* complementing *gid1c*, together with Col-0 wildtype. Seeds were sown on 0.5xMS plates, stratified for 4 days, then grown under diurnal WL for 3 days. Subsequently, seedlings were exposed to either diurnal WL (white) or WL+FR (pink) for 3 days and chemical treatments as indicated. GUS signal accumulation of **(D)** *pGID1A: GID1A-GUS*, **(E)** *pGID1B:GID1B-GUS*, and **(F)** *pGID1C:GID1C-GUS* in 4-day-old seedlings. Red arrows point to differences in protein accumulation and localisation in different light conditions. **(G)** Fragments Per Kilobase of transcript per Million (FPKM) mapped reads of *GID1A, GID1B*, and *GID1C* in Col-0 and *gid1c* hypocotyl + root. **(H)** Emission ratio of YFP/CFP fluorescence in hypocotyls emitted by the GA sensor GPS1. Seedlings were exposed to either diurnal WL or WL+FR for 12 hours before processing for GUS staining (D-F) RNA sequencing (G), or sample fixation and imaging using confocal microscope (H). Data were analysed using two-way (A and C), three-way (B), or one-way ANOVA (H) and Tukey post-hoc analysis. GUS lines were characterised in Suzuki *et al*, 2009.

To examine expression of the three paralogs, we used GID1-GUS reporter lines driven by the endogenous GID1 promoters^14,16^. WL+FR did not visibly alter GID1A or GID1B accumulation or localisation (Figure 1D, E), whereas GID1C accumulated strongly in the hypocotyl and root specifically under WL+FR (Figure 1F). This was not accompanied by a rise in GID1C transcript, which remained low and tended to decrease under WL+FR (Figure 1G), placing the control downstream of transcription. To test whether local bioactive GA level also increased, we imaged hypocotyls of seedlings carrying the FRET-based *GIBBERELLIN PERCEPTION SENSOR 1* (*GPS1*)^17^. No significant change in GA was detected after 12 hours of WL+FR, although exogenous GA was properly reported by GPS1 (Figure 1G). Together, these results indicate that under WL+FR, plants modulate GA sensitivity rather than GA level over this timeframe.

### GID1C is essential for transcriptional reprogramming in response to WL+FR

Transcriptional reprogramming is central to the NP response^18^. Building on our finding that GID1C is required for the GA-dependent NP response and its localisation within plants (Figure 1), we asked which transcriptional changes underlie this requirement. We performed RNA-seq on hypocotyl-root of from Col-0 and *gid1c* seedlings harvested after 12 hours of WL+FR (Figure S2A), and fitted a single factorial DESeq2 model^19,20^ (∼ genotype + light + genotype:light) across four genotypes × light combinations (n = 4 each; 23,909 genes detected). Unlike separate pairwise comparisons, this one factorial model gives every contrast a common dispersion estimate and a consistent sample set, making the light responses of the two genotypes directly comparable. From this model we extracted the light response within each genotype and the genotype difference, and classified genes (adjusted p < 0.01, |log2FC| ≥ 0.8) as GID1C-independent (similar light response in both genotypes), GID1C-affected (light response differs between genotypes), or constitutive genotype (genotype effect with no light response in either genotype).

Of 1,726 DEGs, 1,489 (86.3%) were GID1C-affected, responding to WL+FR differently in Col-0 and *gid1c* (Figure 2A). In contrast, GID1C-independent and constitutive genotype represented a minor fraction. The volcano plots further show that the strong response to WL+FR observed in Col-0 (Figure 2B) is largely lost in *gid1c* (Figure 2C). Within the GID1C-affected set, 1,362 out of 1,489 genes (91.5%) responded in Col-0 and lost significance in *gid1c* (GID1C-required; Figure 2D). Of these, 726 were induced and 636 repressed. 124 (8.3%) genes gained a response in *gid1c*, suggesting GID1C normally inhibit its expression. Next, we plotted the GID1C-affected genes based on their differential expression under WL+FR versus WL in Col-0 (x-axis) and *gid1c* (y-axis; Figure 2E). GID1C-required genes (n = 1362) scattered along the Col-0 axis, responding to WL+FR in Col-0 but remaining near zero in *gid1c*, whereas the smaller set gained in *gid1c* (n = 124) spread along the *gid1c* axis. GID1C is therefore required for the majority of the WL+FR transcriptional response, in both the induced and repressed directions.

**Figure 2.**
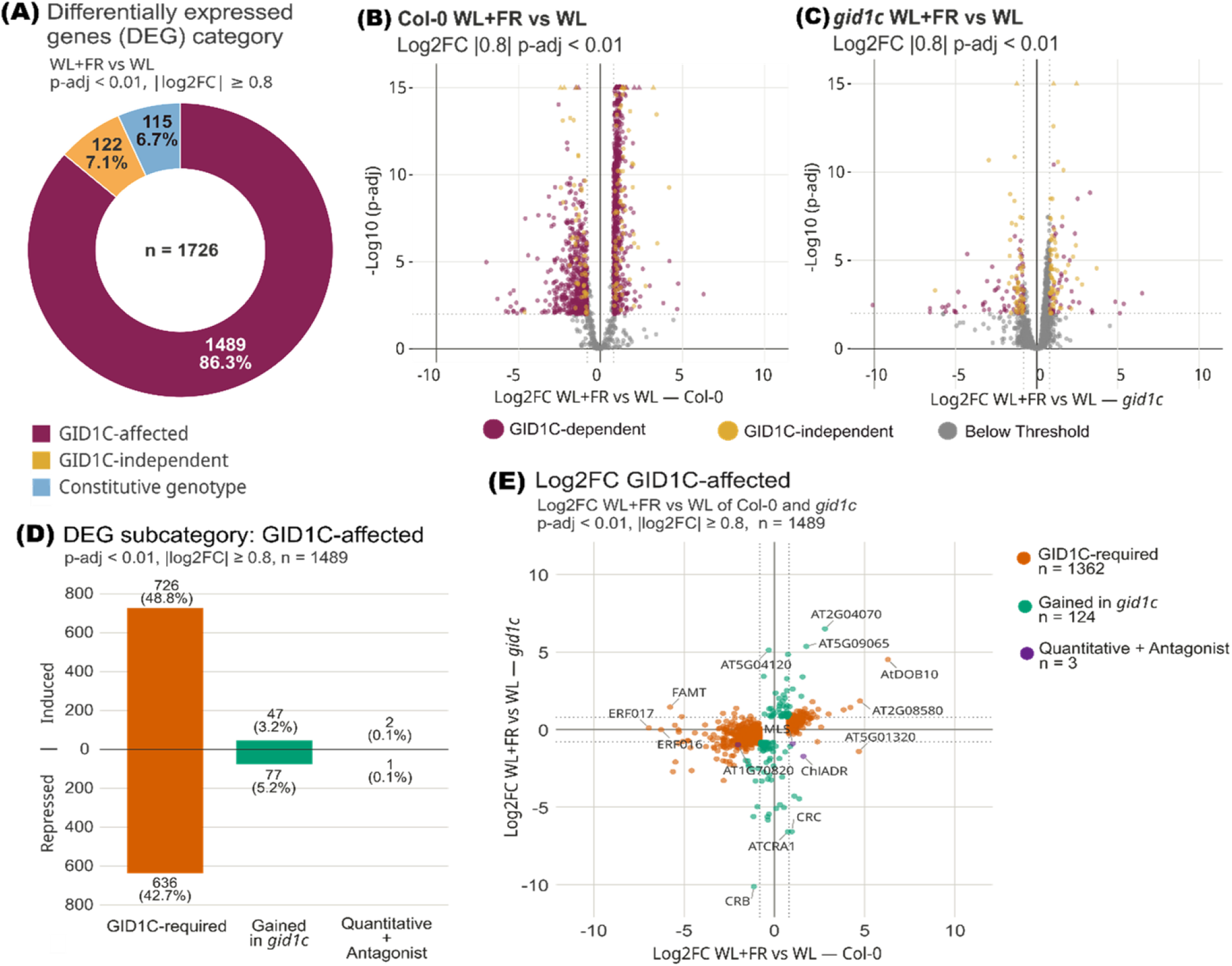
GID1C mediates hypocotyl and root transcriptional reprogramming in neighbour proximity. **(A)** Distribution of DEGs WL+FR vs WL among 3 main groups: GID1C-affected (violet), GID1C-independent (yellow), and light-independent constitutive genotype effect (blue). DEGs classified from a factorial DESeq2 model (∼ genotype + light + genotype:light) using within-genotype simple-effect and genotype main-effect contrasts (p-adj < 0.01, |log2FC| ≥ 0.8). Volcano plot of **(B)** Col-0 wild-type and **(C)** *gid1c* WL+FR vs WL hypocotyl+root, visualising GID1C-dependent (violet), GID1C-independent (yellow), and below-threshold (grey). Thresholds are indicated. **(D)** Barplot and **(E)** scatter plot visualising DEG subgroups within the GID1C-affected group in D, subdivided into GID1C-required (orange), gained in *gid1c* (green), and quantitative+antagonist (purple), with direction indicated. Scatter plot showing log2 fold-change in WL+FR vs WL of Col-0 (x-axis) and *gid1c* (y-axis) with colour-coded genes based on its subgroup in D.

### GID1C acts through transcriptional hubs in the neighbour-proximity response

Gene Ontology analysis of the GID1C-required set revealed GID1C involvement in diverse WL+FR-responsive processes, spanning transcriptional regulation, development and morphogenesis, and stress response, across 35 enriched GO-Biological Process (GO-BP) terms (Figure 3A). Regulation of DNA-templated transcription showed the strongest enrichment (Kolmogorov–Smirnov test p-value = 8.30 × 10^−5^), comprised 97 genes, 22 induced and 75 repressed (Figure 3B). To identify common members across the enriched terms, we counted genes shared by two or more terms. Of 1362 GID1C-required genes, 311 were annotated to more than one enriched GO-BP term. Nine of the ten most recurrent genes were transcription factors and members of the most significantly enriched term (Figure 3C), suggesting that GID1C acts through transcriptional hubs within the WL+FR response. Among these, the third most recurring gene *ICE1* was significantly repressed under WL+FR in Col-0 (log2FC = −0.82, padj = 0.007) but not in *gid1c* (log2FC = −0.50, padj = 0.26; Figure 3B), indicating *ICE1* is repressed under WL+FR in a GID1C-dependent manner. We therefore examined *ICE1* further as a representative node of the GID1C-required programme.

**Figure 3.**
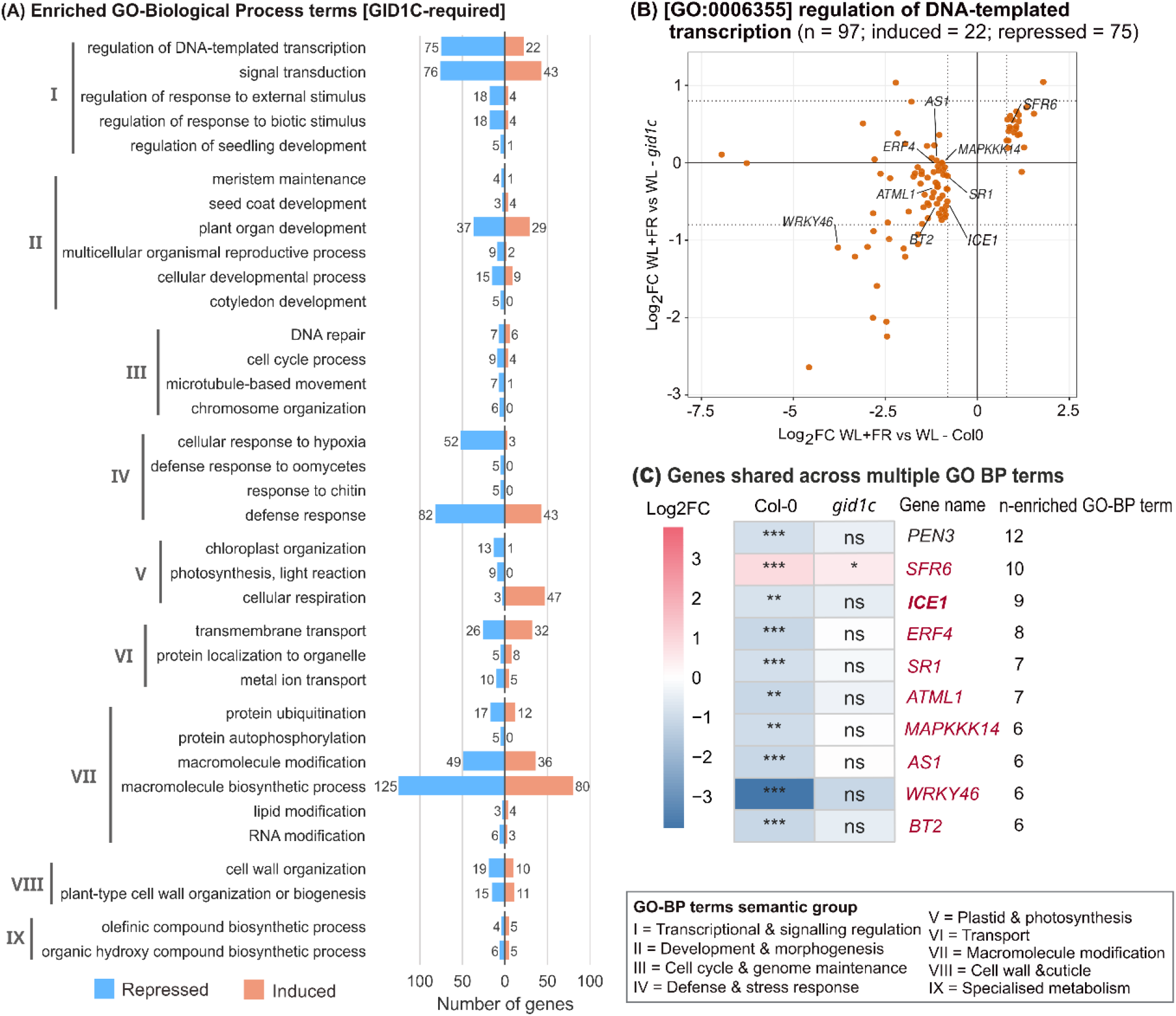
Gene Ontology analysis reveals GID1C involvement in diverse biological processes in response to neighbour proximity. **(A)** Enriched GO-BP terms in GID1C-required gene set (n=1362). GO Term was analysed using Kolmogorov-Smirnov test weight01 algorithm (p<0.01). The number of induced (coral) and repressed (blue) genes in each term are indicated, and grouped by term semantics listed in black box. **(B)** Scatter plot of genes that are significantly enriched in GO-BP Regulation of DNA-templated transcription (GO:0006355). Genes are plotted based on value of log2 fold-change in WL+FR vs WL of Col-0 (x-axis) and *gid1c* (y-axis). **(C)** Log2FC heatmap of Top 10 most shared genes across GO-BP terms with number of enriched GO-BP terms are indicated. Red font indicates its membership in B. Bold font indicates the selected gene of interest.

### A phosphorylation-resistant *ice1* allele disrupts the WL+FR elongation response

We examined *ICE1*, a bHLH transcription factor, as a representative GID1C-required regulatory hub. We measured hypocotyl length across light conditions and GA treatments in two *ICE1* alleles. *ice1-CR*/*scrm-cr*, a CRISPR allele in which a 1665 bp deletion removes almost the entire ICE1 coding sequence^21^, and *ice1-1*, which carries the dominant R236H substitution that prevents MITOGEN-ACTIVATED PROTEIN KINASE 3/6 (MPK3/6)-dependent phosphorylation of ICE1^22^ (Figure 4). *ice1-CR* elongated under WL+FR to the same extent as Col-0 (Figure 4A). Because WL+FR represses *ICE1* (Figure 3C), *ice1-CR* deletion reproduces this repression, resulting in wild-type-like phenotype of *ice1-CR*. In contrast, *ice1-1* failed to elongate under WL+FR and showed identical growth to Col-0 under WL (Figure 4B). The *ice1-1* phenotype therefore reflects an activity introduced by the phosphorylation-resistant R236H substitution, indicating that phosphorylation, alongside transcript repression, may keep ICE1 from inhibiting the WL+FR response. In darkness, *ice1-1* still elongated substantially, although less than Col-0 (Figure 4C), indicating that the phosphorylation-resistant R236H *ice1-1* specifically affects the WL+FR elongation response. To test the GA-responsiveness of *ice1-1*, we pharmacologically manipulated GA levels (Figure 4D). In Col-0, PAC abolished the WL+FR response and exogenous GA_4+7_ enhanced elongation under both light conditions. In contrast, *ice1-1* did not elongate in response to either WL+FR or changes in GA level, indicating insensitivity to both light and GA treatments. ICE1’s inhibition of the WL+FR elongation response is therefore regulated at two levels; (1) transcriptionally through GID1C-dependent repression that lowers *ICE1* abundance, and (2) post-translationally through phosphorylation^22^.

**Figure 4.**
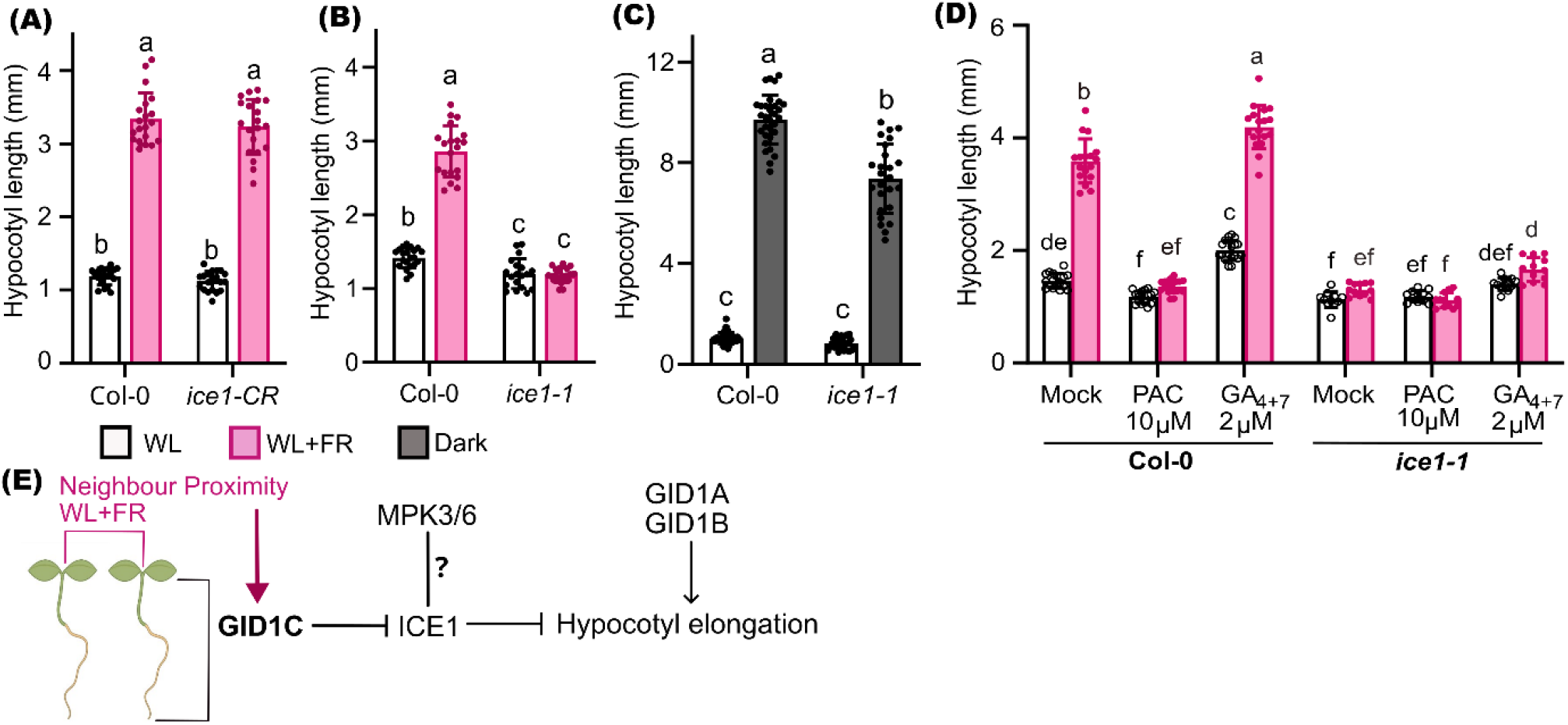
ICE1 is required for full hypocotyl elongation in neighbour proximity. Hypocotyl length of **(A)** Col-0 and *ice1-CR*, **(B)** Col-0 and *ice1-1* in WL+FR (pink) and WL (white). Hypocotyl elongation of Col-0 and *ice1-1* in **(C)** WL and dark (grey), and **(D)** WL+FR and WL in combination with PAC or GA_4+7_. **(E)** Proposed model for GID1C in the NP response. Neighbour proximity (low R:FR, WL+FR) specifically promotes GID1C accumulation, which represses ICE1. ICE1 is proposed to act as a brake on hypocotyl elongation, so that its repression by GID1C promotes elongation. GID1A and GID1B contribute to elongation independently of the light signal, with their accumulation unchanged under WL+FR. The R236H substitution in *ice1-1* blocks MPK3/6-dependent phosphorylation of ICE1. whether this phosphorylation activates or inhibits ICE1 in NP context is unresolved, marked as question mark (?). Seeds were sown on 0.5xMS plates and stratified for 4 days. For A and C, seeds then grown under diurnal WL for 3 days. Subsequently, seedlings were exposed to either diurnal WL or WL+FR and a combination of chemicals as indicated for 3 days. For B, seeds were exposed to WL for 8 hours, then were moved to either dark condition or stay in WL, to grow for 5 days. Data were analysed using two-way (A-B) or three-way (C) ANOVA and Tukey post-hoc analysis.

## Discussion

### GA perception is an active regulatory layer in response to neighbour proximity

Developmental plasticity in plant hormone signalling is facilitated by receptor paralog redundancy, multifunctionality, and cross-talk mechanisms, enabling adaptive responses to variable environments^23,24^. In line with this, our data show that GID1C is the main GA receptor modulating the NP response in hypocotyl-root, increasing GA sensitivity (Figure 1A-C). GID1C represents a previously unrecognised mode of receptor specialisation. Its protein accumulates specifically under WL+FR in the hypocotyl and root tip, without a corresponding rise in transcript or in bioactive GA (Figure 1F–H), giving it a context-specific role that emerges only when the light environment changes. These results point to GA sensitivity as a contributing variable in the WL+FR response, alongside the established roles of GA metabolism. Within the GA pathway, previous work has focused on biosynthesis, catabolism, and DELLA stability ^25–27^. Our data add a further layer of regulation at the level of perception, where receptor identity shapes how the signal is perceived in the responding tissues. While GID1A and GID1B are also expressed in the hypocotyl and contribute to elongation (Figure 1A, D, E), GID1C is the main determinant of an optimal response to neighbour proximity.

### GID1C drives the neighbour-proximity transcriptional response

The key role of GID1C is also reflected by the transcriptomic data. We modelled the RNA-seq data with a DESeq2 generalised linear model^19,20^ (2×2 factorial, genotype × light, extracting the genotype contrast (*gid1c* vs Col-0) and the within-genotype light responses (WL+FR vs WL in each genotype) from a single fit. Because all samples are analysed together, every comparison draws on one pooled estimate of variability rather than its own, placing the two genotypes’ responses on the same scale. Most differentially expressed genes responding to WL+FR in Col-0 but showing little or no response in *gid1c* (Figure 2). This loss of transcriptional response in *gid1c* corresponds to its reduced hypocotyl elongation under low R:FR (Figure 1). GA-bound GID1 receptors promote DELLA degradation to release PIF-mediated transcription^28^, acting downstream of GA perception. Our data place the WL+FR response at GID1C itself, through regulation of receptor abundance without a detectable change in bioactive GA. Loss of GID1C removes most of the low R:FR transcriptional and growth response, identifying GID1C as the primary receptor driving this programme. The residual elongation likely reflects partial redundancy with the other GID1 paralogs, which are expressed in the hypocotyl and contribute to elongation (Figure 1A, D, E), together with GA-independent regulation of DELLA stability^25^ and other signalling inputs that drive elongation independently of GID1C^29–31^. Because GID1C sets the GA sensitivity this response requires, its absence leaves most of the WL+FR transcriptional reprogramming in the hypocotyl-root unexecuted.

### ICE1 acts as a GID1C-repressed brake on the neighbour-proximity response

ICE1/SCRM is a broadly expressed basic helix-loop-helix (bHLH) transcription factor that acts across several developmental and stress contexts. It was first identified as an activator of cold-induced C-REPEAT/DRE BINDING FACTOR (CBF)/DRE BINDING PROTEIN 1 (DREB1) genes and freezing tolerance^32,33^. It was later shown to specify successive steps of the stomatal lineage^34^, myrosin-cell differentiation^35^, and modulates Abscisic Acid responses during germination^36^. Across these contexts, ICE1 is regulated post-translationally, not by transcription alone. OPEN STOMATA 1 (OST1) stabilises ICE1^37^, whereas MPK3/6 phosphorylation destabilises it^38^. Our data identify ICE1 as a regulatory node in the NP response that is controlled at two levels: (1) transcriptionally, through GID1C-dependent repression that lowers ICE1 abundance (Figure 3B), and (2) post-translationally, through phosphorylation (Figure 4). The consequences of MPK3/6 phosphorylation are context-dependent, destabilising ICE1 in the stomatal lineage^22^ while promoting its activity in the embryo^21^. The elongation response inhibition in *ice1-1* is similar with the mechanism seen in stomatal development where phosphorylation inhibits ICE1 activity. However, which route operates under WL+FR remains to be determined.

Together, these findings position GA perception as an active and regulated step of NP response. Rather than passively receiving bioactive GA, the GID1C receptor itself is regulated by light. Perception thus represents a further regulated layer of the NP response, with GID1C regulates the morphological response to low R:FR, and integrates regulatory nodes such as ICE1 within it. The condition-dependent regulation of a single GA receptor paralogs may reflect the diversification of the receptor family across flowering plants^15^. Most eudicots encode several GID1 paralogues, which could permit subfunctionalisation to particular environmental conditions^15,39,40^ that is not available to single-GA-receptor species such as rice^41,42^. Receptor duplication may therefore allow GA responses to be tuned across tissues and conditions, which could make paralogues advantageous in variable environments.

## Material and methods

### Plant material and initial growth conditions

All experiments in this study were performed using *Arabidopsis thaliana*. The used *Arabidopsis* lines are listed in Supplemental Table T1. Seeds were surface sterilised using ethanol absolute, sown on 0.5xMS square plates (pH 5.8, 1% agar), stratified for 4 days, then moved into a controlled environment growth chamber with a 12-h-light/12-h-dark cycle, temperature of 20°C and a light level of PAR ∼100 μmol/m^2^/s (WL). Seeds were germinating and grown vertically in this condition for the length stated in the specific methods below.

### Hypocotyl length assay

Seeds were germinated under initial growth condition for 3 days. On the 4^th^ day, uniformly grown seedlings were selected and transferred into a fresh 0.5xMS plates. For light treatment, plates contained transferred seedlings were placed vertically into a controlled environment growth chamber with a 12-h-light/12-h-dark cycle of WL, or WL supplemented with additional far-red LED (WL+FR, FR = 57 μmol/m^2^/s). For combination with GA treatment, seedlings were transferred into 0.5xMS plates with 2µM GA_4+7_, 10µM PAC, or solvent (ethanol/mock), before subjecting them with WL or WL+FR. The seedlings were grown in these conditions for 3 days before scanning the plates using Epson V700 double-sided scanner. Hypocotyl length of the seedlings was measured from the image using FIJI/ImageJ.

### GUS staining

*pGID1A:GID1A-GUS, pGID1B:GID1B*, and *pGID1C:GID1C-GUS* seeds were germinated under initial growth condition for 4 days. At ZT0 of the 5^th^ day, seedlings were treated with WL or WL+FR for 12 hours (n>20), before fixing and staining step according to previously described method^43^. Stained seedlings were mounted into microscopy slides, then scanned using Epson V700 double-sided scanner.

### GA sensor and confocal imaging

GPS1 seeds were germinated under initial growth condition for 4 days. At ZT0 of the 5^th^ day, seedlings were treated with WL, WL+FR, or WL+2µM GA_4+7_ for 12 hours (n>20), before fixing and clearing step according to Clearsee-α method^44^. Confocal imaging was performed on Leica SP8 systems equipped with multi-photon lasers, according to previously published method^17,45^. Detailed confocal microscope setup is described in Supplemental Table T2.

### RNA sequencing sample harvesting and processing

Col-0 and *gid1c* seeds were germinated under initial growth condition for 4 days. At ZT0 of the 5th day, seedlings were treated with WL or WL+FR for 12 hours. Next, seedlings were vacuum-infiltrated and incubated in liquid 0.5xMS containing 1mM cordycepin for 15 minutes to inhibit RNA degradation^46^. Seedling were washed with liquid 0.5xMS, dried, and cotyledons were removed from hypocotyl-root (n>80) for each set of genotype and light treatment (n=4), inserted into 1.5 mL tubes, and snap-frozen in liquid nitrogen. Total RNA of hypocotyl+root samples was isolated using RNeasy Plant Mini Kit (Qiagen) and then proceed to poly-A mRNA sequencing.

### RNA sequencing data processing and analysis

Raw gene counts of the hypocotyl+root samples (n=16 libraries) were obtained from the sequencing service provider. Fragments per kilobase per million (FPKM) values were computed for expression-level annotation only and did not enter differential-expression testing. Sample-level quality was assessed by principal component analysis (PCA, Figure S2B-C). For differential expression, genes with fewer than 10 summed reads across libraries were removed, and a negative-binomial generalised linear model (∼ genotype + light + genotype:light) was fit jointly across all 16 libraries using DESeq2 version 3.23 in R^19,47^. From the fitted model we extracted the genotype × light interaction, the main genotype and light contrasts, and the within-genotype simple effects of light in each genotype. Fold changes are reported as maximum-likelihood log2 estimates. Genes were considered responsive at adjusted p-value < 0.01 (Benjamini-Hochberg) and |log2FC| ≥ 0.8, and NP-responsive genes were classified from their simple-effect patterns as GID1C-independent (both genotypes responding similarly in direction and magnitude) or GID1C-dependent (any differential pattern). Threshold choices were supported by sensitivity analyses across a range of log2FC and magnitude-ratio cutoffs (Figure S2D-E). Gene Ontology enrichment analysis was performed using Kolmogorov–Smirnov weight01 algorithm using topGO in R^48^.

### Statistical analysis and data visualisation

Hypocotyl length and GA sensor data were analysed on R using ANOVA and Tukey posthoc corresponds to each experimental design. Plot visualisation and figure arrangement was done using ggplot2 R and inkscape, respectively.

## Acknowledgements

The authors thank Dr. Patrick Achard (University of Strasbourg) and Prof. Masatoshi Nakajima (University of Tokyo) for GID1-GUS lines, and Dr. Alexander Jones (Sainsbury Lab) for the GA sensor GPS1 lines. The authors also thank Prof. Marcel Quint, Dr. Carolin Delker, and Dr. Kasper van Gelderen for hosting some of the experiments reported here.

## References

1. Ballaré, C. L. & Pierik, R. The shade-avoidance syndrome: multiple signals and ecological consequences: Neighbour detection and shade avoidance. Plant, Cell & Environment 40, 2530–2543 (2017).

2. Casal, J. J. Photoreceptor Signaling Networks in Plant Responses to Shade. Annual Review of Plant Biology 64, 403–427 (2013).

3. van Gelderen, K. et al. Gibberellin transport affects lateral root growth through HY5 in response to far-red light. Plant Cell 37, koaf200 (2025).

4. Galvao, V. C., Horrer, D., Kuttner, F. & Schmid, M. Spatial control of flowering by DELLA proteins in Arabidopsis thaliana. Development 139, 4072–4082 (2012).

5. Bou-Torrent, J. et al. Plant proximity perception dynamically modulates hormone levels and sensitivity in Arabidopsis. Journal of Experimental Botany 65, 2937–2947 (2014).

6. Lopez-Juez, E., Kobayashi, M., Sakuraia, A., Kamiya, Y. & Kendrick, R. E. Phytochrome, Gibberellins, and Hypocotyl Growth. Plant Physiology 107, 131–140 (1995).

7. Weller, J. L., Ross, J. J. & Reid, J. B. Gibberellins and phytochrome regulation of stem elongation in pea. Planta 192, 489–496 (1994).

8. Islam, S., Park, K., Xia, J., Kwon, E. & Kim, D. Y. Structural insights of gibberellin-mediated DELLA protein degradation. Molecular Plant S1674205225002035 (2025) doi:10.1016/j.molp.2025.06.010.

9. Ueguchi-Tanaka, M., Nakajima, M., Motoyuki, A. & Matsuoka, M. Gibberellin Receptor and Its Role in Gibberellin Signaling in Plants. Annual Review of Plant Biology 58, 183–198 (2007).

10. Iuchi, S. et al. Multiple loss-of-function of Arabidopsis gibberellin receptor AtGID1s completely shuts down a gibberellin signal: Multiple knockout mutant for gibberellin receptors. The Plant Journal 50, 958–966 (2007).

11. Griffiths, J. et al. Genetic Characterization and Functional Analysis of the GID1 Gibberellin Receptors in Arabidopsis. the Plant Cell Online 18, 3399–3414 (2006).

12. Prasetyaningrum, P., Mariotti, L., Valeri, M. C. & Novi, G. Nocturnal gibberellin biosynthesis is carbon dependent and adjusts leaf expansion rates to variable conditions. Plant Physiology (2020).

13. Rieu, I. et al. Genetic Analysis Reveals That C19-GA 2-Oxidation Is a Major Gibberellin Inactivation Pathway in Arabidopsis. the Plant Cell Online 20, 2420–2436 (2008).

14. Suzuki, H. et al. Differential expression and affinities of Arabidopsis gibberellin receptors can explain variation in phenotypes of multiple knock-out mutants. The Plant Journal 60, 48–55 (2009).

15. Yoshida, H. et al. Evolution and diversification of the plant gibberellin receptor GID1. Proceedings of the National Academy of Sciences 201806040 (2018) doi:10.1073/pnas.1806040115.

16. Shi, B. et al. A quantitative gibberellin signaling biosensor reveals a role for gibberellins in internode specification at the shoot apical meristem. Nat Commun 15, 3895 (2024).

17. Rizza, A., Walia, A., Lanquar, V., Frommer, W. B. & Jones, A. M. In vivo gibberellin gradients visualized in rapidly elongating tissues. Nature Plants 3, 803–813 (2017).

18. Kohnen, M. V. et al. Neighbor detection induces organ-specific transcriptomes, revealing patterns underlying hypocotyl-specific growth. Plant Cell 28, 2889–2904 (2016).

19. Love, M. I., Huber, W. & Anders, S. Moderated estimation of fold change and dispersion for RNA-seq data with DESeq2. Genome Biol 15, 550 (2014).

20. Law, C. W. et al. A guide to creating design matrices for gene expression experiments. Preprint at 10.12688/f1000research.27893.1 (2020).

21. Chen, H. et al. Phosphorylation-dependent activation of the bHLH transcription factor ICE1/SCRM promotes polarization of the Arabidopsis zygote. New Phytologist 245, 1029–1039 (2025).

22. Putarjunan, A. et al. Bipartite anchoring of SCREAM enforces stomatal initiation by coupling MAP kinases to SPEECHLESS. Nat. Plants 5, 742–754 (2019).

23. Spartz, A. K. & Gray, W. M. Plant hormone receptors: new perceptions. Genes Dev 22, 2139–2148 (2008).

24. Wong, C., Alabadí, D. & Blázquez, M. A. Spatial regulation of plant hormone action. J Exp Bot 74, 6089–6103 (2023).

25. Blanco-Touriñán, N. et al. COP1 destabilizes DELLA proteins in Arabidopsis. Proceedings of the National Academy of Sciences of the United States of America 117, 13792–13799 (2020).

26. Djakovic-Petrovic, T., Wit, M. D., Voesenek, L. A. C. J. & Pierik, R. DELLA protein function in growth responses to canopy signals. Plant Journal 51, 117–126 (2007).

27. Küpers, J. J. et al. Local light signaling at the leaf tip drives remote differential petiole growth through auxin-gibberellin dynamics. Current Biology 33, 75–85.e5 (2023).

28. Feng, S. et al. Coordinated regulation of Arabidopsis thaliana development by light and gibberellins. Nature 451, 475–479 (2008).

29. Hayes, S., Velanis, C. N., Jenkins, G. I. & Franklin, K. A. UV-B detected by the UVR8 photoreceptor antagonizes auxin signaling and plant shade avoidance. Proc. Natl. Acad. Sci. U.S.A. 111, 11894–11899 (2014).

30. Keuskamp, D. H. et al. Blue-light-mediated shade avoidance requires combined auxin and brassinosteroid action in Arabidopsis seedlings. Plant Journal 67, 208–217 (2011).

31. Michaud, O., Fiorucci, A.-S., Xenarios, I. & Fankhauser, C. Local auxin production underlies a spatially restricted neighbor-detection response in Arabidopsis. Proc. Natl. Acad. Sci. U.S.A. 114, 7444–7449 (2017).

32. Chinnusamy, V. et al. ICE1: a regulator of cold-induced transcriptome and freezing tolerance in Arabidopsis. Genes Dev. 17, 1043–1054 (2003).

33. Tang, K. et al. The transcription factor ICE1 functions in cold stress response by binding to the promoters of CBF and COR genes. Journal of Integrative Plant Biology 62, 258–263 (2020).

34. Kanaoka, M. M. et al. SCREAM/ICE1 and SCREAM2 Specify Three Cell-State Transitional Steps Leading to Arabidopsis Stomatal Differentiation. Plant Cell 20, 1775–1785 (2008).

35. Shirakawa, M. et al. FAMA Is an Essential Component for the Differentiation of Two Distinct Cell Types, Myrosin Cells and Guard Cells, in Arabidopsis. Plant Cell 26, 4039–4052 (2014).

36. Hu, Y. et al. The Transcription Factor INDUCER OF CBF EXPRESSION1 Interacts with ABSCISIC ACID INSENSITIVE5 and DELLA Proteins to Fine-Tune Abscisic Acid Signaling during Seed Germination in Arabidopsis. Plant Cell 31, 1520–1538 (2019).

37. Ding, Y. et al. OST1 Kinase Modulates Freezing Tolerance by Enhancing ICE1 Stability in Arabidopsis. Developmental Cell 32, 278–289 (2015).

38. Li, H. et al. MPK3- and MPK6-Mediated ICE1 Phosphorylation Negatively Regulates ICE1 Stability and Freezing Tolerance in Arabidopsis. Developmental Cell 43, 630–642.e4 (2017).

39. Xu, P. et al. Blue light-dependent interactions of CRY1 with GID1 and DELLA proteins regulate gibberellin signaling and photomorphogenesis in Arabidopsis. The Plant Cell 33, 2375–2394 (2021).

40. Mjema, E. Y., Bonatelli, M. L., Consortium, T. N. & Laubinger, S. Molecular and phenotypic footprints of climate in native Arabidopsis thaliana. 2026.03.02.709013 Preprint at 10.64898/2026.03.02.709013 (2026).

41. Ueguchi-Tanaka, M. et al. Molecular Interactions of a Soluble Gibberellin Receptor, GID1, with a Rice DELLA Protein, SLR1, and Gibberellin. The Plant Cell 19, 2140–2155 (2007).

42. Gazara, R. K., Moharana, K. C., Bellieny-Rabelo, D. & Venancio, T. M. Expansion and diversification of the gibberellin receptor GIBBERELLIN INSENSITIVE DWARF1 (GID1) family in land plants. Plant Mol Biol 97, 435–449 (2018).

43. Malamy, J. E. & Benfey, P. N. Organization and cell differentiation in lateral roots of Arabidopsis thaliana. Development 124, 33–44 (1997).

44. Kurihara, D., Mizuta, Y., Nagahara, S. & Higashiyama, T. ClearSeeAlpha: Advanced Optical Clearing for Whole-Plant Imaging. Plant Cell Physiol 62, 1302–1310 (2021).

45. Rizza, A. et al. Differential biosynthesis and cellular permeability explain longitudinal gibberellin gradients in growing roots. 10.1073/pnas.1921960118 (2021) doi:10.1073/pnas.1921960118.

46. Prasetyaningrum, P. et al. Inhibition of RNA degradation integrates the metabolic signals induced by osmotic stress into the Arabidopsis circadian system. Journal of Experimental Botany erad274 (2023) doi:10.1093/jxb/erad274.

47. DESeq2. Bioconductor http://bioconductor.org/packages/DESeq2/.

48. topGO. Bioconductor http://bioconductor.org/packages/topGO/.

