## Supplemental data for "Functional specialization of the gibberellin receptor GIBBERELLIN-INSENSITIVE DWARF 1C in plant neighbour detection"

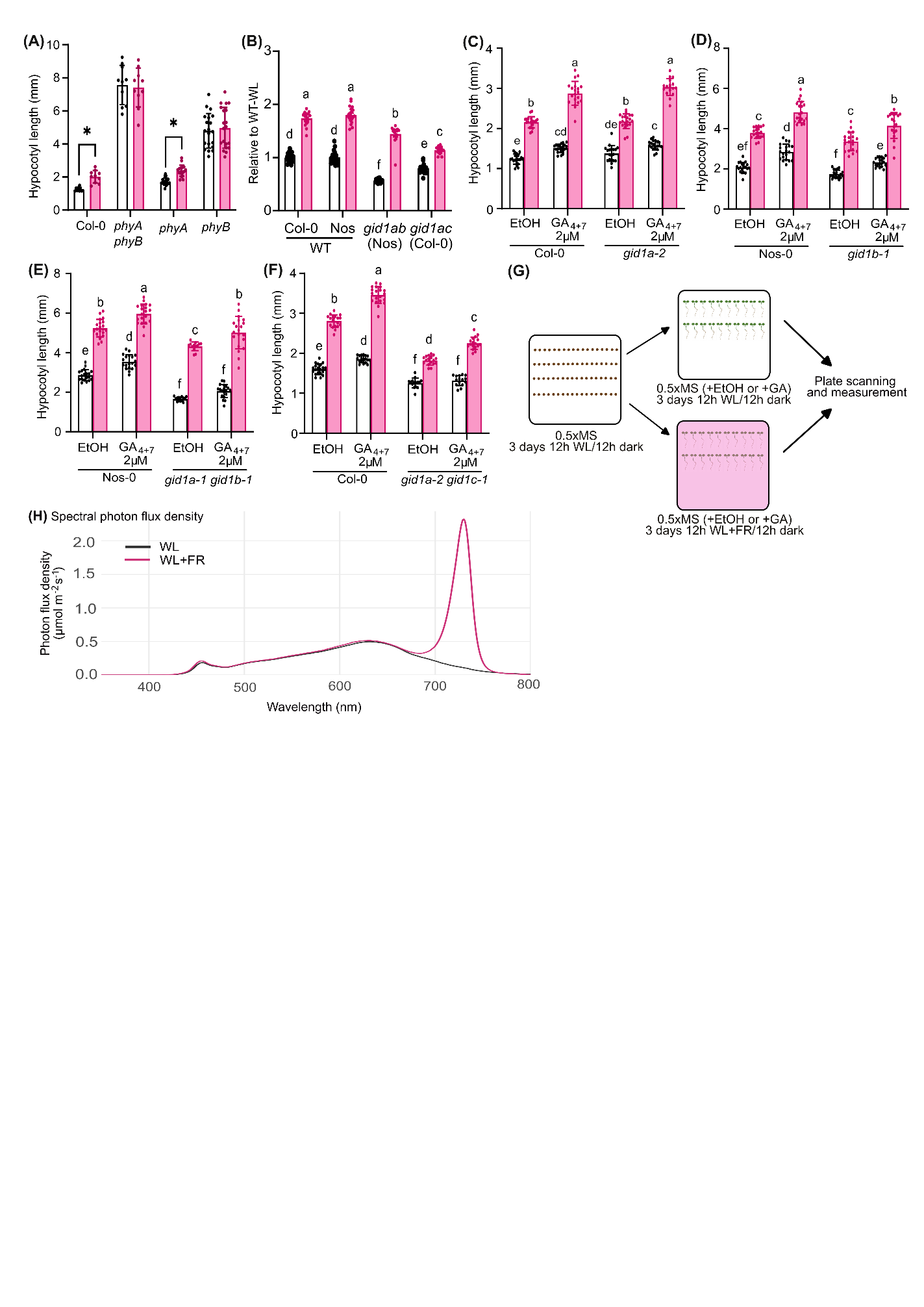
**SUPPLEMENTAL FIGURES**

**Supplemental Figure S1.** Hypocotyl length of 7 days old **(A)** Col-0, *phya-211, phyb-9,* and *phya-211 phyb-9*, **(B)** *gid1ab* (Nos) and *gid1ac* (Col-0) and their respective wildtype, **(C)** *gid1a-1* and its wildtype, **(D)** *gid1b-1* and its wildtype, **(E)** *gid1a-1 gid1b-1* double mutant and its wildtype, **(F)** *gid1a-2 gid1c-1* and its wildtype, after 3 days of WL+FR (pink) treatment and WL (white) control condition. **(G)** Experimental procedure and **(H)** Spectral photon flux density of WL and WL+FR used in this entire report. seedlings were exposed to either diurnal WL (white) or WL+FR (pink) for 3 days (A-F) and a combination of chemicals as indicated (B-F). Data were analysed using two-way (A ), or three-way (B-F) and Tukey post-hoc analysis for main light effect (A) or interaction effect (B-F)

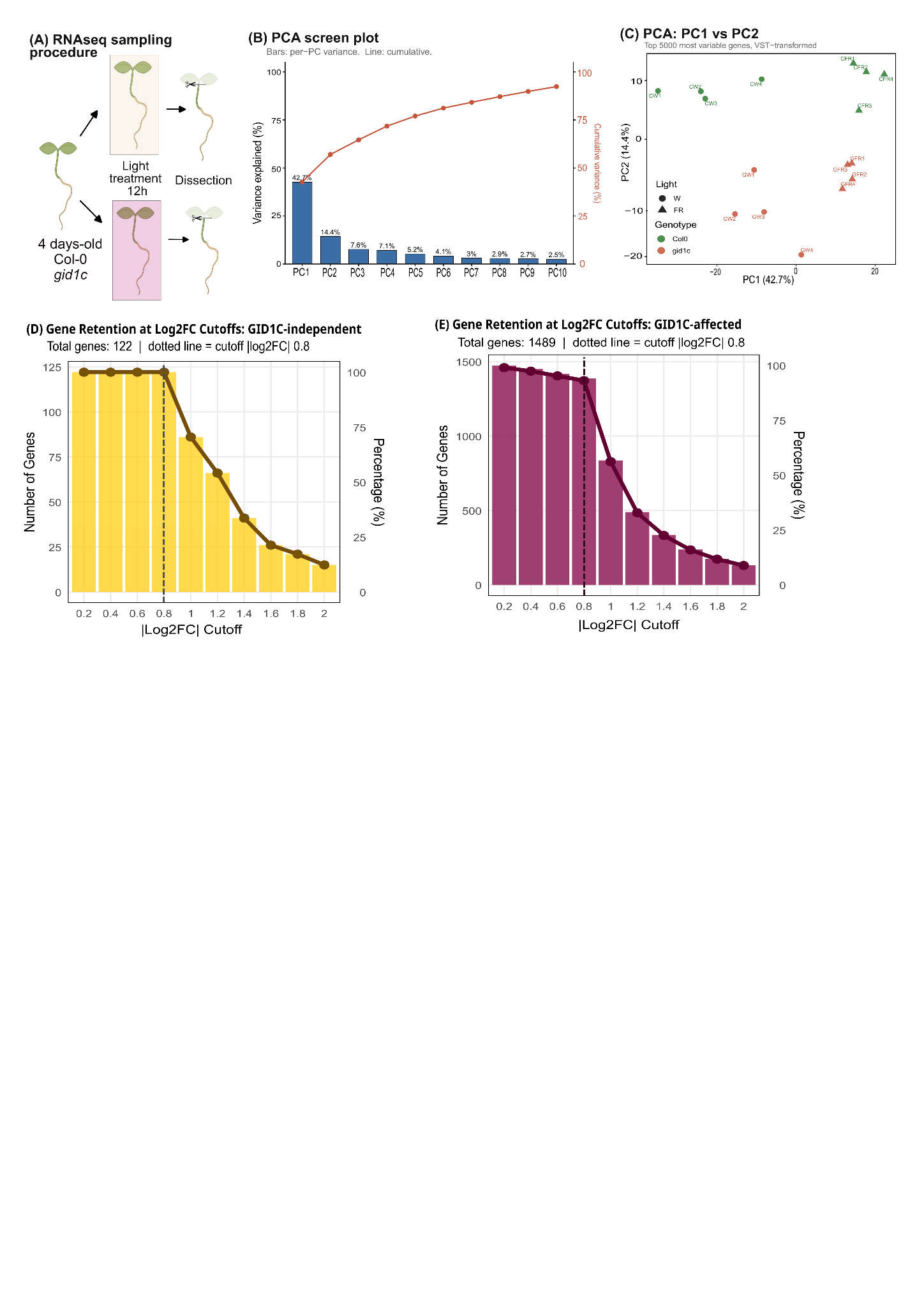

**Supplemental Figure S2.** **(A)** Sampling scheme. Four-day-old Col-0 (wild type) and *gid1c* seedlings were treated for 12 h with either WL or WL+FR to simulate neighbour proximity. Hypocotyl+root of the seedlings were then separated from their cotyledons and processed for RNA sequencing. **(B)** Bar-and-line plot of the principal component analysis. Bars give the variance explained by each component (PC1–PC10) and the line gives the cumulative variance; PC1 and PC2 together account for 57.1%. **(C)** Sample ordination on PC1 (42.7%) versus PC2 (14.4%), computed from the 5,000 most variable genes after variance-stabilising transformation (VST). Point shape denotes light treatment (circle, W; triangle, FR) and colour denotes genotype (green for Col-0, orange for *gid1c*); each point is one biological replicate. **(D, E)** Sensitivity of gene number to the fold-change threshold applied to the Col-0 light response, shown for the **(D)** GID1C-independent (n = 122) and **(E)** GID1C-affected (n = 1,489) gene sets. Bars show the number of genes retained at each |log2FC| cutoff (left axis) and the line show the corresponding percentage of the set (right axis). The dashed vertical line marks the |log2FC| ≥ 0.8 membership threshold (applied with Padj < 0.01).

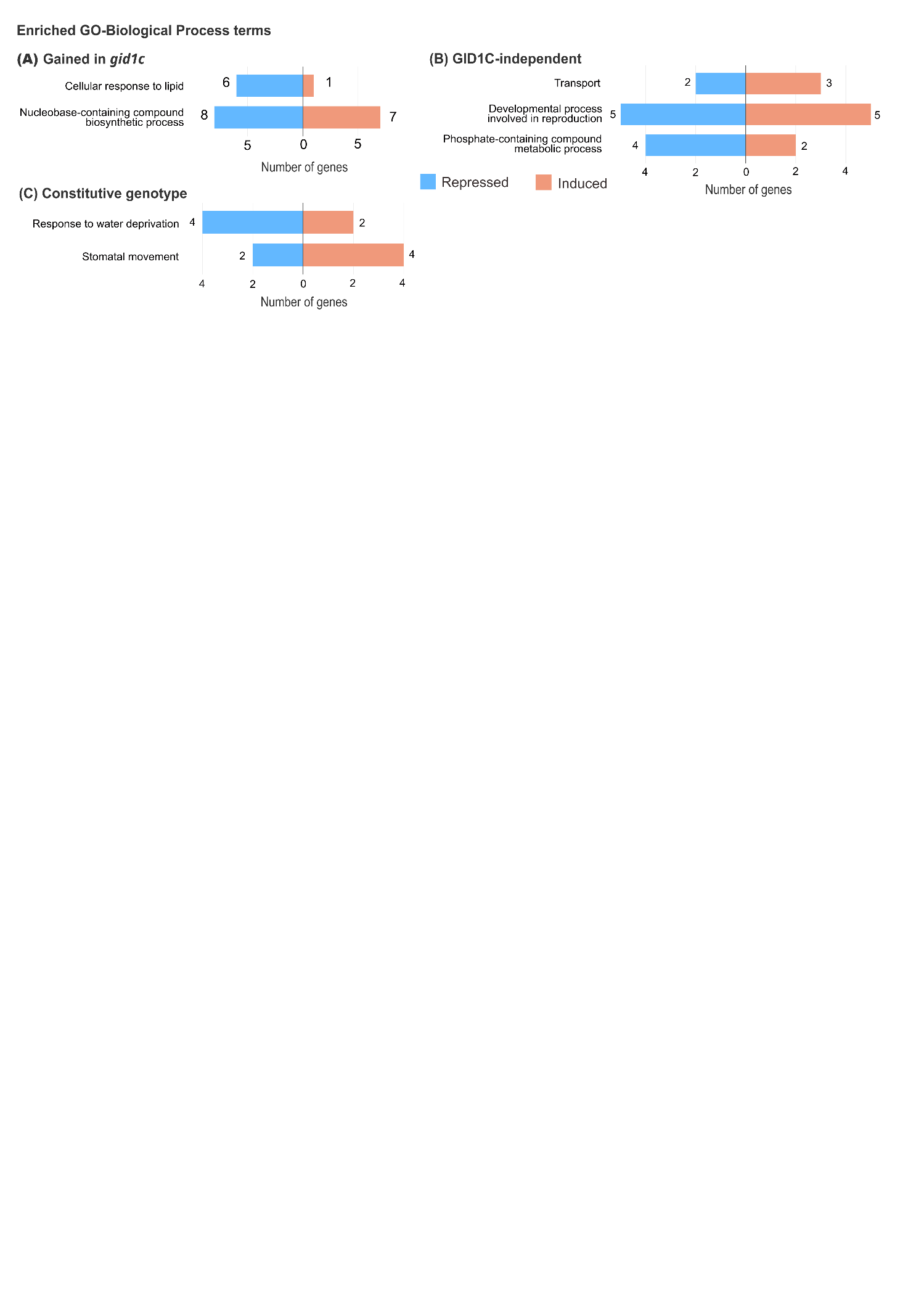

**Supplemental Figure S3.** Diverging bar plots of significantly enriched Gene Ontology biological-process (GO BP) terms of DEGs for three categories of shade-responsive genes: **(A)** Genes gaining a response in gid1c (Gained in *gid1c*), **(B)** GID1C-independent genes, and **(C)** constitutive genotype-difference genes. For each term, bars show the number of genes moving in each direction under low R:FR: repressed to the left (blue) and induced to the right (coral), with the gene count labelled at each bar end. The vertical line marks zero. Terms were called as enriched using the topGO Kolmogorov–Smirnov test (weight01 algorithm).

| **Genotype** | **Background** | **Reference** |
| --- | --- | --- |
| Columbia-0 (Col-0/wildtype) | N/A | N/A |
| Nossen-0 (Nos/wildtype, NASC ID: N1394) | N/A | Iuchi et al., 2007; Suzuki et al., 2009; Yoshida et al., 2018 |
| *gid1a-2* | Col-0 |  |
| *gid1b-1* | Nos |  |
| *gid1c-1* | Col-0 |  |
| *gid1a-1 gid1b-1* | Nos |  |
| *gid1a-2 gid1c-1* | Col-0 |  |
| *pGID1C:GID1C-mCitrine* | *gid1c* | This study |
| *pGID1A:GID1A-GUS* | Col-0 | Shi et al., 2024; Suzuki et al., 2009 |
| *pGID1B:GID1B-GUS* | Col-0 |  |
| *pGID1B:GID1B-GUS* | Col-0 |  |
| *GPS1* | Col-0 | (Rizza et al., 2017) |
| *ice1-1* | Col-0 | Kidokoro et al., 2020 |
| *ice1-CR* | Col-0 | Chen et al., 2025 |
| *phyA-211* | Col-0 | Viczián et al., 2020 |
| *phyB-9* | Col-0 |  |
| *phyA-211 phyb-9* | Col-0 |  |

**Supplemental Table T1. *Arabidopsis* lines used in this study**

**Supplemental Table T2. Confocal microscope setup**

| **Confocal settings** | | | | | | |
| --- | --- | --- | --- | --- | --- | --- |
| **Name** | | | | **Value** | | |
| Scan Mode | | | | xyz | | |
| Scan Direction X | | | | Bidirectional | | |
| Scan Speed | | | | 200 Hz | | |
| Magnification | | | | 20 | | |
| ObjectiveName | | | | HC PL APO CS2 20x/0.75 IMM | | |
| Immersion | | | | IMM | | |
| Numerical Aperture | | | | 0.75 | | |
| RefractionIndex | | | | 1.518 | | |
| Zoom | | | | 2.4 | | |
| **Lasers** | | | | | | |
| WLL | | | | On, 85.00 %, Maximum Power | | |
| **Setting 1: YFP-control (For nuclei segmentation)** | | | | | | |
| **Device Name** | | | | **Value** | | |
| Pinhole | | | | 77.6 µm | | |
| PinholeAiry | | | | 1.5 AU | | |
| EmissionWavelength for PinholeAiry Calculation | | | | 530 nm | | |
| **Laser Line** | | | | **Intensity** | | |
| Laser Line (405 nm) | | | | Intensity: 0.00% | | |
| Laser Line (449 nm) | | | | Intensity: 0.00% | | |
| Laser Line (514 nm) | | | | Intensity: 5.00% | | |
| Laser Line (442 nm) | | | | Intensity: 0.00% | | |
| Laser Line (443 nm) | | | | Intensity: 0.00% | | |
| Laser Line (787 nm) | | | | Intensity: 0.00% | | |
| Laser Line (788 nm) | | | | Intensity: 0.00% | | |
| Laser Line (789 nm) | | | | Intensity: 0.00% | | |
| Laser Line (790 nm) | | | | Intensity: 0.00% | | |
| **Detectors** | | | | | | |
| **Detector Name** | **Operating Mode** | **Reference Line** | **Spectral Positions / Gain / Offset** | | **Image Channel** | **Channel Information** |
| HyD S 1 | Analog | --- | (435nm - 440nm) / 2.5 / na | | --- | --- |
| HyD X 2 | Digital | --- | (456nm - 500nm) / 10 / na | | --- | --- |
| HyD S 3 | Counting | --- | (500nm - 506nm) / na / na | | --- | --- |
| HyD R 4 | Digital | 514 | (520nm - 588nm) / 244.1 / na | | Intensity | --- |
| Trans PMT | Analog |  | n/a / 23.5 / 0 | | Analog | --- |
| **Setting 2: CFP_Ex – YFP_Em detection (FRET signal)** | | | | | | |
| **Device Name** | | | | **Value** | | |
| Pinhole | | | | 77.6 µm | | |
| PinholeAiry | | | | 1.5 AU | | |
| EmissionWavelength for PinholeAiry Calculation | | | | 530 nm | | |
| **Laser Line** | | | | **Intensity** | | |
| Laser Line (405 nm) | | | | Intensity: 0.00% | | |
| Laser Line (449 nm) | | | | Intensity: 0.00% | | |
| Laser Line (514 nm) | | | | Intensity: 5.00% | | |
| Laser Line (442 nm) | | | | Intensity: 0.00% | | |
| Laser Line (443 nm) | | | | Intensity: 0.00% | | |
| Laser Line (787 nm) | | | | Intensity: 0.00% | | |
| Laser Line (788 nm) | | | | Intensity: 0.00% | | |
| Laser Line (789 nm) | | | | Intensity: 0.00% | | |
| Laser Line (790 nm) | | | | Intensity: 0.00% | | |
| **Detectors** | | | | | | |
| **Detector Name** | **Operating Mode** | **Reference Line** | **Spectral Positions / Gain / Offset** | | **Image Channel** | **Channel Information** |
| HyD S 1 | Analog | --- | (435nm - 440nm) / 300 / na | | --- | --- |
| HyD X 2 | Digital | 449 | (456nm - 500nm) / 160 / na | | Intensity | --- |
| HyD S 3 | Counting | --- | (500nm - 506nm) / na / na | | --- | --- |
| HyD R 4 | Digital | 449 | (520nm - 588nm) / 450 / na | | Intensity | --- |
| Trans PMT | Analog | --- | / 0 / 0 | | --- | --- |
